# Deletion of HindIIR and HindIIIR improves DNA transfer via electroporation to *Haemophilus influenzae* Rd

**DOI:** 10.1101/2024.07.09.602704

**Authors:** Samir Hamadache, Yu Kang Huang, Adam Shedeed, Aqil Syed, Bogumil J. Karas

**Affiliations:** Department of Biochemistry, Schulich School of Medicine and Dentistry, The University of Western Ontario, London, ON N6A 5C1, Canada

**Keywords:** *Haemophilus influenzae*, genetic toolbox, multi-host shuttle plasmids, restriction-modification, electroporation

## Abstract

*Haemophilus influenzae* is a bacterial species of interest for its medical relevance and utility as a model system. Despite its role in several landmark molecular and synthetic biology studies, *H. influenzae* remains underexplored as a potential chassis organism. The limited availability of reliable and convenient transformation methods and genetic tools for *H. influenzae* are obstacles to this end. However, a strain of *H. influenzae* Rd KW20 lacking the type II restriction endonucleases HindII and HindIII has previously been developed. Here, we show that this strain is more readily transformable by electroporation than wild-type Rd KW20. We also developed a series of multi-host plasmids carrying antibiotic selection and fluorescent visual markers based on the pSU20 vector. The availability of *H. influenzae* ΔHindII/III, paired with the electroporation method and plasmids presented here, will promote the exploration of *H. influenzae* as a host organism for synthetic biology applications.

## 1. INTRODUCTION

*Haemophilus influenzae* is found in two places: in humans, where it is generally commensal in the respiratory and genital tracts, and in the laboratory [1]. The species was first described in 1892 by Richard Pfeiffer, who believed it to be the cause of influenza [2]. Since then, *H. influenzae* has been implicated in several milestones of molecular biology and studied for its medical relevance.

*H. influenzae* is a gram-negative, pleomorphic, facultatively anaerobic coccobacillus that requires hemin and nicotinamide adenine dinucleotide (NAD) for aerobic growth [3]. Sizing only 0.3–1.0 µm, it has a short doubling time of ∼30 minutes and displays natural competence for DNA uptake, an interesting area of study in its own right [4,5]. *H. influenzae* strain Rd KW20 is the first bacterium to have its complete genome sequenced [6]. Its small, circular genome (1.8 Mbp with 38% GC content) has raised interest among synthetic biologists in creating a new minimal cell, with its entire genome having already been cloned in *S. cerevisiae* [7]. Nevertheless, *H. influenzae* remains underexplored as a chassis, owing to a dearth of genetic tools and reliable transformation methods.

Strains of *H. influenzae* have been categorized according to their various capsular polysaccharide antigens from types a through f, with unencapsulated strains deemed nontypeable [8]. *H. influenzae* Rd is derived from a type d strain selected for its loss of encapsulating antigens [9]. Rd KW20 was later derived from this strain and used in several studies by Hamilton Smith and colleagues, including the Nobel prize-winning discovery of the first type II restriction enzyme, HindII [10,11]. Rd KW20 is non-pathogenic, although it has virulence properties, allowing for its use as a model system for invasive *H. influenzae* [12].

The common mode of transforming *H. influenzae* is the MIV method, named after the nongrowth starvation media (chemically defined medium M-IV) that induces *H. influenzae*’s natural competence [13]. This method requires linear DNA containing a 9-bp uptake signal sequence [14,15]. Since competent *H. influenzae* is highly proficient at homologous recombination, this approach is practical for genome insertions where the transforming DNA contains sufficient homology to the chromosome. However, using the MIV method to transform plasmids is inefficient; plasmids can be linearized for uptake, but without sufficient homology for recombination, they will quickly be degraded by exonucleolytic activity [14]. *H. influenzae* is transformable by other methods; however, these methods have not always been reproducible [16–19].

According to REBASE, *H. influenzae* Rd encodes up to eight restriction-modification (RM) systems in its genome (**Supplementary Figure S1**), two of which (HindII and HindIII) are well-characterized type II restriction systems [20]. Bacteria possess RM systems primarily as innate immune protection against bacteriophages, although other functions of these systems have also been explored [21]. In 2013, Karas et al. deleted the HindII and HindIII restriction endonucleases in the Rd KW20 genome, improving its delivery to yeast cells [7]. Here, we show that the “ΔHindII/III” strain generated by Karas et al. can reliably be transformed by electroporation when using plasmids containing HindII and/or HindIII restriction sites. We also present a series of plasmids based on the pSU20 shuttle vector carrying various selectable and visual markers.

## 2. RESULTS AND DISCUSSION

### 2.1 *H. influenzae* ΔHindII/III transforms at a higher efficiency compared to wild-type

*H. influenzae* ΔHindII/III was developed by Karas et al. by replacing HindIIR with a cassette containing tetracycline and ampicillin resistance genes and HindIIIR with a chloramphenicol resistance gene [7]. While these markers were initially used to create the deletions, they provide limited antibiotic resistance (**Supplementary Figure S2**). We sought to determine whether this strain could be transformed more reliably with pSU20 and related plasmids compared to the wild-type strain using electroporation. To do this, we developed an electroporation protocol for *H. influenzae* and used it to deliver plasmids conferring chloramphenicol resistance to each strain (see methods). Furthermore, it is also possible to use tetracycline or kanamycin as selection (**Supplementary Figure S3**).

Overall, ΔHindII/III consistently yielded greater numbers of transformants as compared with wild-type *H. influenzae* (**Figure 1A**) and individual ΔHindII/III transformant colonies screened by restriction digestion confirmed the presence of plasmids with no gross rearrangements (**Figure 1B**). All ΔHindII/III transformants tested produced the expected banding pattern whereas for WT, 1/10 pSU20 transformants (#5, **Figure 1B**) and 2/10 pHflu4 transformants (#7 and #9, **Figure 1B**) did not yield the expected digest products (for additional analysis see **Supplementary Figure S4**). Interestingly while wild-type *H. influenzae* did produce some transformant colonies, substantial background growth/biofilm was observed on these plates as early as 24 hours after incubation, including those where sterile water was instead of DNA used as negative controls (**Supplementary Figure S5**). This background growth was reduced when smaller volumes of recovered cells were plated (**Supplementary Figure S5**). Future work investigating this background growth phenotype and its absence on ΔHindII/III transformation plates may be of interest.

**Figure 1.**
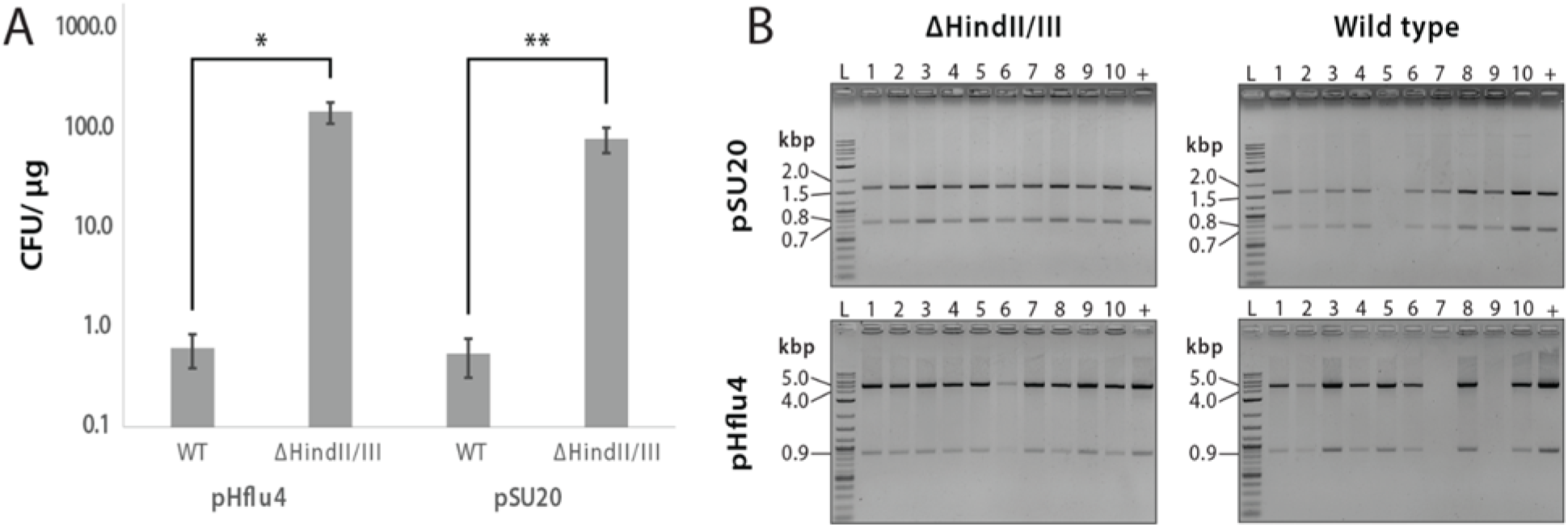
*H. influenzae* ΔHindII/III transforms more efficiently by electroporation compared to wild-type. (**A**) Transformation competent cells were prepared from wild-type (“WT”, n = 4) and ΔHindII/III (n = 4) and then electroporated with either pSU20 or pHflu4 plasmid DNA. ΔHindII/III produced more colony forming units (CFU) per microgram of transforming DNA (*: p = 0.12, **: p = 0.04). Error bars represent standard error from the mean. (**B**) Restriction digest of plasmids isolated from ΔHindII/III and WT transformants (+: original plasmid used for transformation, L: NEB 1kb plus ladder)

The effect of restriction endonucleases on bacterial transformation efficiency is well-understood and it has been shown in several species that the removal of restriction systems improves transformation [22]. pSU20 contains HindII and HindIII recognition sites within its multiple cloning region and pHflu4 contains five HindIII and two HindII recognition sites. These recognition sites are fairly prevalent across commonly used vector plasmids such as pUC19 as they are typically included in multiple cloning sites. Our work suggests that other plasmids carrying HindII or HindIII recognition sequences could also be used in *H. influenzae* ΔHindII/III. The availability of an *H. influenzae* Rd KW20 variant that is more readily transformed by electroporation should accelerate research in this species and promote its development as a chassis for synthetic biology.

### 2.2 Selectable and fluorescent markers for *H. influenzae*

The most used vector plasmids are the pSU series (mainly pSU20), derived from pACYC184, which provides moderate copy numbers in *Escherichia coli* and *H. influenzae* [23]. The presence of HindII and HindIII recognition sites is less problematic when *H. influenzae* is transformed by the MIV method; DNA is taken up as single strands in this approach, whereas only double-stranded DNA acts a substrate for restriction endonucleases [14]. With ΔHindII/III and the convenience of electroporation in hand, we generated a series of plasmids carrying fluorescent and selectable markers for *H. influenzae* (**Figure 2**).

**Figure 2.**
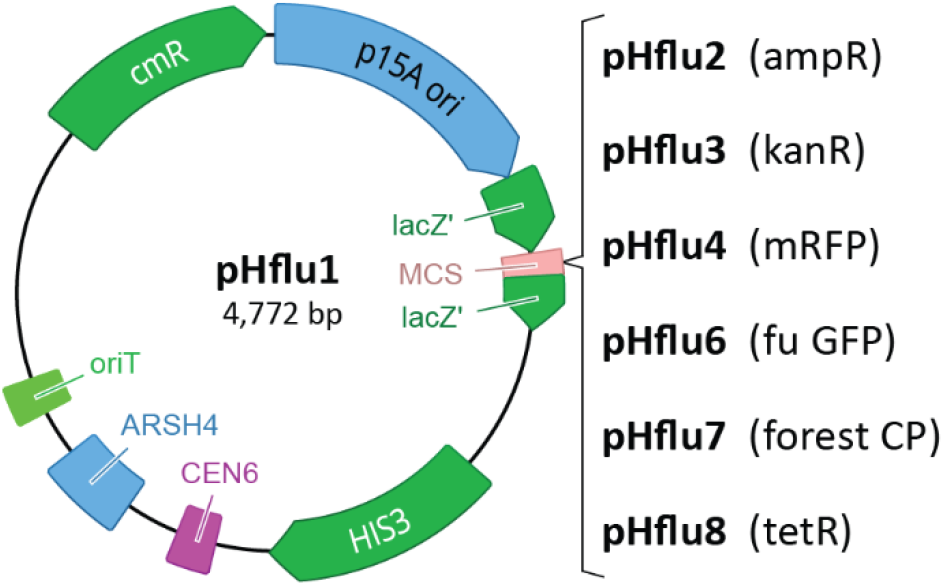
Plasmid map of pHflu1 indicating where markers were inserted to generate pHflu2-8.

We first assembled pHflu1 by adding elements to pSU20 for plasmid maintenance in *S. cerevisiae* (CEN6, ARSH4, HIS3) and the RP4/RK2 origin of conjugal transfer (oriT). The former was added to enable assembly of pHflu1-based plasmids in yeast, and the latter to enable the possibility of delivering plasmids to *H. influenzae* by bacterial conjugation. In addition to the chloramphenicol acetyltransferase (which provides resistance to chloramphenicol) present on pHflu1, we then added an ampicillin resistance gene (*bla*) to make pHflu2, one for kanamycin (*nptII*), making pHflu3, and one for tetracycline (*tetM*), giving pHflu8. Upon successfully electroporating these plasmids into ΔHindII/III, we spot-plated serial dilutions of the resulting transformants alongside ΔHindII/III carrying no plasmid or pHflu1 onto plates containing the appropriate antibiotics (**Figure 3**). This screen confirmed the antibiotic resistance conferred by the appropriate plasmids.

**Figure 3.**
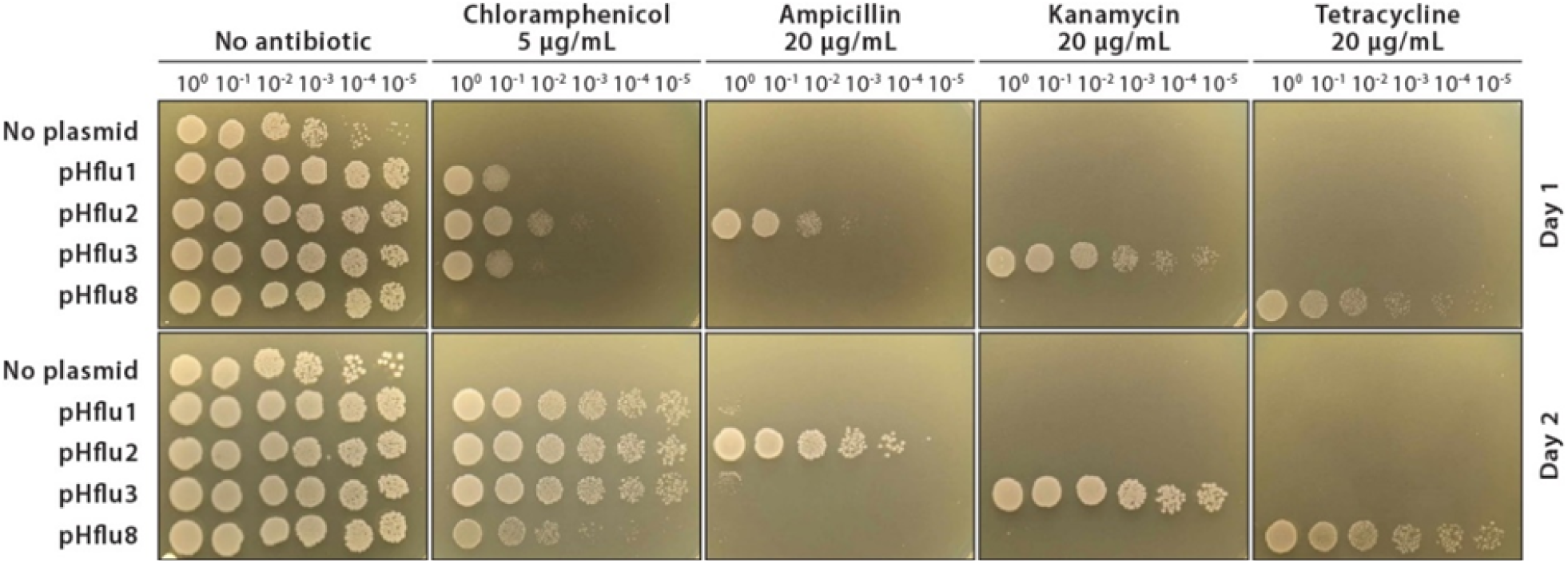
pHflu2, pHflu3, and pHflu8 confer resistance to ampicillin, kanamycin, and tetracycline respectively, in *H. influenzae* ΔHindII/III. Serial dilutions of *H. influenzae* ΔHindII/III transformants harboring either no plasmid, pHflu1, pHflu2, pHflu3, or pHflu8 were plated on sBHI supplemented with antibiotics as indicated. Images were taken after one and two days of growth at 37 °C.

We then generated three variants of pHflu1 carrying fluorescent visual markers. Monomeric RFP (mRFP) was inserted to make pHflu4 and two additional fluorophores obtained from a collaborator were inserted, making pHfu6 (fu GFP) and pHflu7 (forest CP). pHflu4 is visibly red when plated on sBHI media containing 5 µg/mL of chloramphenicol, whereas pHflu6 and pHflu7 appear green under ultraviolet light. All three fluorescent markers are clearly visible by fluorescence microscopy (**Figure 4**).

**Figure 4.**
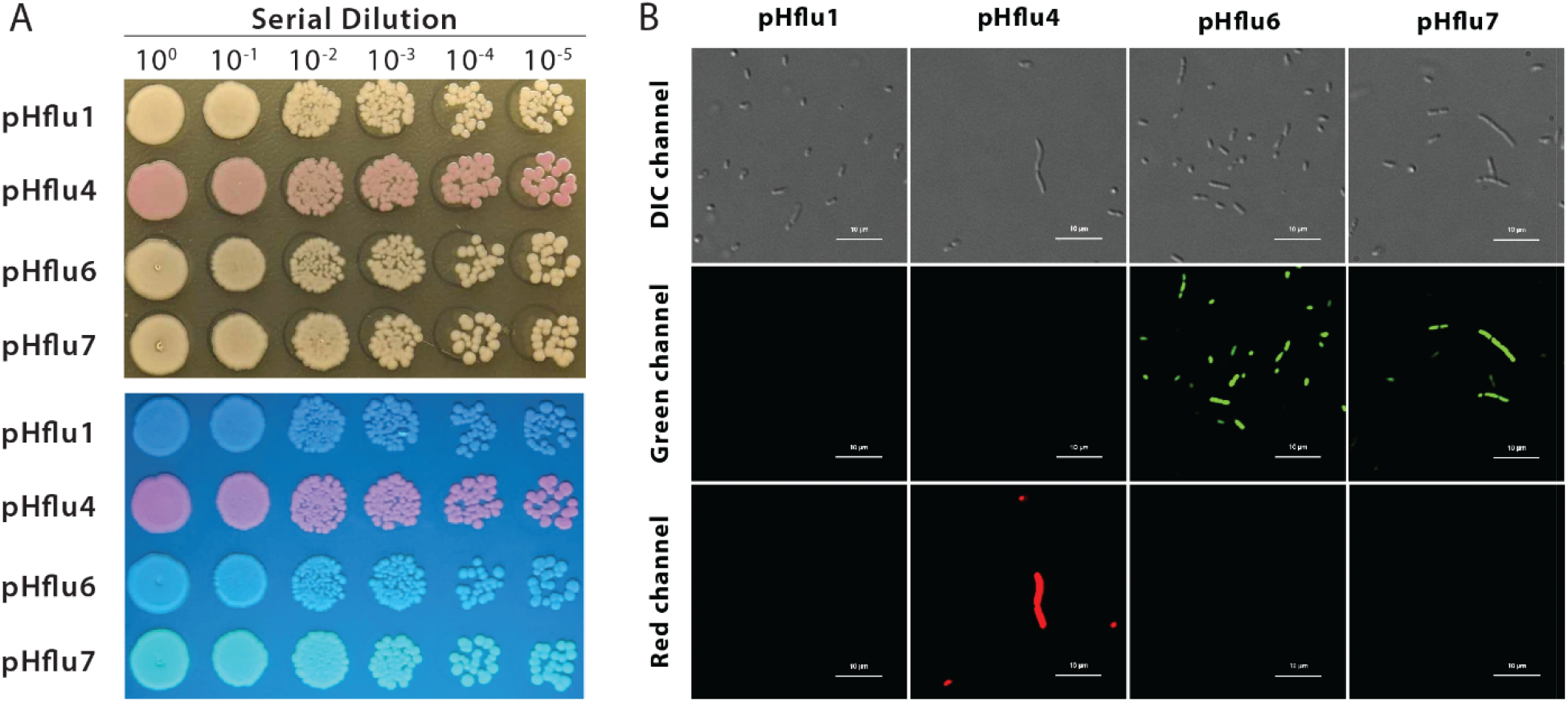
Visualization of pHflu1 variants carrying visual markers. (**A**) Red fluorescence provided by pHflu4 is visible in ambient light, while green provided by pHflu6 and pHflu7 is better observed under UV light. *H. influenzae* ΔHindII/III transformants were serially diluted and spot-plated on sBHI supplemented with 5 µg/ml chloramphenicol and grown for 2 days at 37 °C. The image below was taken under UV light. (**B**) *H. influenzae* ΔHindII/III carrying pHflu1, pHflu4, pHflu6, and pHflu7 as observed by confocal fluorescence microscopy. Red channel excitation/emission wavelength: 594/613 nm; red channel: 488/543 nm

## Conclusions

*H. influenzae* lacking HindII and HindIII endonuclease genes yields a greater number of colonies than Rd KW20 when electroporated with plasmids containing recognition sites for these restriction enzymes. Additionally, we present a series of plasmids based on the pSU20 multi-host shuttle vector carrying different selection and visual markers. Future work could examine if ΔHindII/III similarly improves plasmid transformation by other methods, such as chemical transformation. In addition, the pHflu plasmid series could be used to evolve ΔHindII/III for greater electroporation efficiency as has been done for *Acholeplasma laidlawii* [24]. The electroporation method and genetic tools herein will promote exploration of *H. influenzae* as a synthetic biology chassis organism.

## 3. MATERIALS AND METHODS

### 3.1 Microbial strains and growth conditions

*H. influenzae* Rd KW20 (ATCC 51907) and *H. influenzae* ΔHindII/III [7] were grown in brain and hearth infusion broth (BD Difco™ 237500) supplemented with 20 mg/L of hemin and 4 g/L of NAD (“sBHI”), prepared as described in [25]. MAX Efficiency™ *E. coli* DH5α (Invitrogen™ 18258012) and TransforMax™EPI300™ *E. coli* (LGC Biosearch Technologies EC300110) were grown in LB medium (10 g/L tryptone, 5 g/L yeast extract, 10 g/L NaCl). Both bacterial species were grown at 37 °C with shaking at 225 rpm; for solid media, *E. coli* was grown on 1.5% agar plates, while *H. influenzae* was grown on 1.2% agar plates. For antibiotic selection, LB was supplemented with 15 µg/mL of chloramphenicol, and sBHI was supplemented with 3–5 µg/mL of chloramphenicol, 20 µg/mL of ampicillin, or 20 µg/mL of kanamycin, as appropriate.

*S. cerevisiae* VL6-48 (ATCC MYA-3666) was grown in 2×YPAD medium (20 g/L yeast extract, 40 g/L peptone, 40 g/L glucose, 160 mg/L adenine hemisulfate). Transformed spheroplasts were grown in complete minimal glucose media lacking histidine (Teknova C7112) supplemented with 1 M D-sorbitol as described in [25]. Yeast was grown at 30 °C, shaking at 225 rpm or on 2% agar plates.

### 3.2 Assembly of pHflu1 in *S. cerevisiae*

All primer sequences are provided in the **Supplementary Information**.

pHflu1 was constructed via yeast spheroplast transformation as described previously [26]. A cassette containing a centromere (CEN6), autonomous replicating sequence (ARSH4), and histidine complementation marker (HIS3) for *S. cerevisiae*, as well as the RK2/RP4 origin of transfer sequence (oriT) were amplified by PCR from pTA-MOB 2.0 [26] using primers SH01–02. pSU20 [23] was linearized by PCR using primers SH03– 04. The resulting amplicons were purified, combined, and mixed with *S. cerevisiae* spheroplasts, yielding the circular plasmid pHflu1. DNA was isolated from pooled yeast transformants and electroporated, according to manufacturer’s instructions into TransforMax™ EPI300™ *E. coli* where clones were screened by multiplex PCR for validation (using primers SH05–08). pHflu1 is available at https://www.addgene.org/ ID: 203878.

### 3.3 Assembly of pHflu1 variants in *E. coli*

pHflu2–4 and 6–8 were assembled via *E. coli* chemical transformation as previously described [27]. For pHflu2, 3, and 6–8, pHflu1 was linearized by PCR at its multiple cloning site using primers SH09–10. *bla* (beta lactamase) was PCR-amplified from pUC19 using primers SH11–12 to afford pHflu2. *nptII* (aminoglycoside phosphotransferase) was PCR-amplified from pAGE3.0 using primers SH13–14 to afford pHflu3. forest CP and fuGFP were obtained by PCR-amplifying fluorophores from plasmids a gift from Binomica Labs (binomicalabs.org) using primers SH19–20 to afford pHflu6–7. *tetM*, amplified from pAL1 using SH21-22, was likewise inserted into linearized pHflu1 to create pHflu8 [24]. pHflu4 was assembled using pHflu1 linearized by primers SH15–16 and mRFP amplified by primers SH17–18 from a plasmid obtained from Dr. Thomas Hamilton. pHflu2-4 and pHflu6-7 are available at https://www.addgene.org/ IDs: 203880, 203881, 203882, 214087, and 214088 respectively.

### 3.4 Preparation of electrocompetent *H. influenzae*

Single colonies of *H. influenzae* Rd KW20 WT and ΔHindII/III were used to inoculate 3 mL of Difco sBHI in sterile 15-mL conical centrifuge tubes and incubated overnight at 37°C with shaking at 225 rpm. In the morning, the cultures were diluted to an OD_600_ of 0.01 into 25 mL of room temperature sBHI in 125-mL culture flasks and grown for approximately 4 hours at 37 °C with shaking at 225 rpm to reach an OD_600_ of 0.55–0.65 was reached. 4 hours at 37°C with shaking at 225 rpm. The resulting cultures were centrifuged at 3,197 rcf in sterile 50-mL conical centrifuge tubes at 4 °C for 10 minutes and the supernatants decanted. The pelleted cells were resuspended by gentle pipetting in about 12.5 mL of sterile water (kept at 4 °C). This centrifugation and resuspension step was repeated twice more. After a final centrifugation (3,197 rcf, 4 °C, 10 minutes), the supernatant was completely removed, and the pelleted cells were resuspended in 1/10th the original culture volume (250 µL) of 10% glycerol (kept at 4 °C). The cells were then divided into 50 µl aliquots in sterile microcentrifuge tubes, frozen in a -80 °C ethanol bath, and stored at -80 °C for future use.

### 3.5 Electroporation of H. influenzae

50 µL of electrocompetent *H. influenzae* prepared as described above were kept on ice along with 1-mm electrocuvettes and transforming DNA. Transforming DNA was isolated from *E. coli* cultures via alkaline lysis and isopropanol precipitation as previously described [28]. 5 μL of undiluted DNA (sterile water for negative controls) was added to the cells and mixed by gently pipetting twice. The cell-DNA mixture was transferred to a chilled electrocuvette and pulsed using a Gene Pulser Xcell Electroporation System (Bio-Rad) set to 1800 V, 200 Ω, and 50 μF. Time constants obtained averaged around 10.0 ms. 1 mL of room temperature sBHI was immediately added to the electrocuvette and mixed by pipetting. The resulting cell suspension was transferred to a microcentrifuge tube and incubated at 37 °C with 225 rpm shaking for 90 minutes. 1 μL of the recovered cells (mixed with 150 μL of sBHI) were spread onto sBHI plates containing 1.2% agar and the appropriate antibiotic(s) (3 µg/mL of chloramphenicol for pSU20, pHflu1, pHflu4, and pHflu6-7; 20 µg/mL of ampicillin for pHflu2; 20 µg/mL of kanamycin for pHflu3; and 5 μg/mL of tetracycline for pHflu8). For transformations requiring more colonies, 10–100 μL of recovered cells were plated. Colonies were counted after 36-48 hours’ incubation at 37 °C.

### 3.6 Plasmid isolation & restriction digestion assay

Plasmids were isolated from *E. coli* and *H. influenzae* cultures using the EZ-10 DNA Miniprep Kit (BioBasic Inc. BS414) according to the manufacturer’s instructions. DNA was isolated from *S. cerevisiae* cultures via alkaline lysis and isopropanol precipitation as previously described [28]. Isolated pSU20 and pHflu4 were digested using ApoI-HF (New England Biolabs #R3566) and EcoRI-HF (New England Biolabs # R3101), respectively. ∼0.5 ng of plasmid DNA was combined with 2.5 µL of rCutSmart™ (New England Biolabs #B6004), 5 units of enzyme, and deionized water for a total reaction volume of 25 µL. Reaction mixtures were incubated at 37 °C for 15 minutes, followed by 20 minutes at 80 °C to denature the enzymes before separating reaction products on a 1.4% agarose gel by electrophoresis.

### 3.7 Statistical analysis

To compare transformation efficiencies of the WT and dKO strains, colony counts between strains were compared using a two-tailed Student’s T test assuming unequal variances and α = 0.05.

## AUTHOR CONTRIBUTIONS

Conceptualization: S.H., B.J.K; Methodology: S.H., Y.K.H., A.Sh., A.Sy., B.J.K.; Validation: S.H., Y.K.H., A.Sh., B.J.K.; Formal analysis: S.H., A.Sh.; Investigation: S.H., Y.K.H., A.Sh., A.Sy., B.J.K.; Resources: B.J.K.; Writing – Original draft: S.H., Y.K.H.; Writing – Review and Editing: S.H., Y.K.H., B.J.K.; Visualization: S.H., Y.K.H., B.J.K.; Supervision: S.H., B.J.K.; Project administration: B.J.K.; Funding acquisition: B.J.K.

## ACKNOWLEDGMENT

This work was supported by the Government of Canada’s New Frontiers in Research Fund (NFRF), [NFRFE-2018-01124]. Research in B.J.K. laboratory is also supported by Natural Sciences and Engineering Research Council of Canada (NSERC), [RGPIN-2018-06172]. All *H. influenzae* strains used in this study were provided by The J. Craig Venter Institute.

## Notes

### Competing Interest Statement

The authors have declared no competing interest.

## REFERENCES

[1] J.Z. Jordens, M.P. Slack, Haemophilus influenzae: Then and now, Eur. J. Clin. Microbiol. Infect. Dis. 14 (1995) 935–948. 10.1007/BF01691374.

[2] J.K. Taubenberger, J. V Hultin, D.M. Morens, Discovery and characterization of the 1918 pandemic influenza virus in historical context, Antivir. Ther. 12 (2007) 581–591. 10.1177/135965350701200s02.1.

[3] N.M. Evans, D.D. Smith, A.J. Wicken, Haemin and nicotinamide adenine dinucleotide requirements of Haemophilus influenzae and Haemophilus parainfluenzae, J. Med. Microbiol. 7 (1974) 359–365. 10.1099/00222615-7-3-359.

[4] Z.E. Khattak, F. Anjum, Haemophilus influenzae Infection, StatPearls Publishing, Treasure Island (FL), 2023.

[5] D. Dubnau, DNA uptake in bacteria, Annu. Rev. Microbiol. 53 (1999) 217–244. 10.1146/annurev.micro.53.1.217.

[6] R.D. Fleischmann, M.D. Adams, O. White, R.A. Clayton, E.F. Kirkness, A.R. Kerlavage, C.J. Bult, J.F. Tomb, B.A. Dougherty, J.M. Merrick, K. McKenney, G. Sutton, W. FitzHugh, C. Fields, J.D. Gocayne, J. Scott, R. Shirley, L.I. Liu, A. Glodek, J.M. Kelley, J.F. Weidman, C.A. Phillips, T. Spriggs, E. Hedblom, M.D. Cotton, T.R. Utterback, M.C. Hanna, D.T. Nguyen, D.M. Saudek, R.C. Brandon, L.D. Fine, J.L. Fritchman, J.L. Fuhrmann, N.S.M. Geoghagen, C.L. Gnehm, L.A. McDonald, K. V. Small, C.M. Fraser, H.O. Smith, J.C. Venter, Whole-genome random sequencing and assembly of Haemophilus influenzae Rd, Science. 269 (1995) 496–512. 10.1126/science.7542800.

[7] B.J. Karas, J. Jablanovic, L. Sun, L. Ma, G.M. Goldgof, J. Stam, A. Ramon, M.J. Manary, E.A. Winzeler, J.C. Venter, P.D. Weyman, D.G. Gibson, J.I. Glass, C.A. Hutchison, H.O. Smith, Y. Suzuki, Y. Suzuki, Direct transfer of whole genomes from bacteria to yeast., Nat. Methods. 10 (2013) 410–412. 10.1038/nmeth.2433.

[8] M. Pittman, Variation and type specificity in the bacterial species hemophilus influenzae, J. Exp. Med. 53 (1931) 471–471. 10.1084/jem.53.4.471.

[9] G. Leidy, E. Hahn, H.E. Alexander, In vitro production of new types of hemophilus influenzae., J. Exp. Med. 97 (1953) 467–482. 10.1084/jem.97.4.467.

[10] K.W. Wilcox, H.O. Smith, Isolation and characterization of mutants of Haemophilus influenzae deficient in an adenosine 5’ triphosphate dependent deoxyribonuclease activity, J. Bacteriol. 122 (1975) 443–453. 10.1128/jb.122.2.443-453.1975.

[11] H.O. Smith, K.W. Wilcox, A restriction enzyme from Hemophilus influenzae. I. Purification and general properties., Biotechnology. 51 (1970) 379–391. 10.1016/0022-2836(70)90149-X.

[12] D.A. Daines, L.A. Cohn, H.N. Coleman, K.S. Kim, A.L. Smith, Haemophilus influenzae Rd KW20 has virulence properties, J. Med. Microbiol. 52 (2003) 277–282. 10.1099/jmm.0.05025-0.

[13] R.M. Herriott, E.M. Meyer, M. Vogt, Defined nongrowth media for stage II development of competence in Haemophilus influenzae., J. Bacteriol. 101 (1970) 517–524. 10.1128/jb.101.2.517-524.1970.

[14] G. Poje, R.J. Redfield, Transformation of Haemophilus influenzae., Methods Mol. Med. 71 (2003) 57–70. 10.1385/1-59259-321-6:57.

[15] J.C. Mell, I.M. Hall, R.J. Redfield, Defining the DNA uptake specificity of naturally competent Haemophilus influenzae cells, Nucleic Acids Res. 40 (2012) 8536–8549. 10.1093/nar/gks640.

[16] M.A. Mitchell, K. Skowronek, L. Kauc, S.H. Goodgal, Electroporation of Haemophilus influenzae is effective for transformation of plasmid but not chromosomal DNA, Nucleic Acids Res. 19 (1991) 3625–3628. 10.1093/nar/19.13.3625.

[17] N.K. Notani, J.K. Setlow, D. Mccarthy, N.-L. Clayton, Transformation of Haemophilus influenzae by Plasmid RSF0885, J. Bacteriol. 148 (1981) 812–816.

[18] F.J. Pagotto, H. Salimnia, P. a Totten, J.R. Dillon, Stable shuttle vectors for Neisseria gonorrhoeae, Haemophilus spp. and other bacteria based on a single origin of replication, Gene. 244 (2000) 13–19.

[19] M.S. Chandler, New shuttle vectors for Haemophilus influenzae and Escherichia coli: P15A-derived plasmids replicate in H. influenzae Rd, Plasmid. 25 (1991) 221–224. 10.1016/0147-619X(91)90016-P.

[20] R.J. Roberts, T. Vincze, J. Posfai, D. Macelis, REBASE: a database for DNA restriction and modification: enzymes, genes and genomes, Nucleic Acids Res. 51 (2023). 10.1093/nar/gkac975.

[21] K. Vasu, V. Nagaraja, Diverse Functions of Restriction-Modification Systems in Addition to Cellular Defense, Microbiol. Mol. Biol. Rev. 77 (2013) 53–72. 10.1128/mmbr.00044-12.

[22] D. Chung, J. Farkas, J. Westpheling, Overcoming restriction as a barrier to DNA transformation in Caldicellulosiruptor species results in efficient marker replacement, Biotechnol. Biofuels. 6 (2013). 10.1186/1754-6834-6-82.

[23] B. Bartolomé, Y. Jubete, E. Martínez, F. de la Cruz, Construction and properties of a family of pACYC184-derived cloning vectors compatible with pBR322 and its derivatives, Gene. 102 (1991) 75–78. 10.1016/0378-1119(91)90541-I.

[24] D.P. Nucifora, N.D. Mehta, D.J. Giguere, B.J. Karas, An expanded genetic toolbox to accelerate the creation of Acholeplasma laidlawii driven by synthetic genomes Graphical Abstract, (n.d.). 10.1101/2022.09.21.508766.

[25] B.J. Karas, J. Jablanovic, E. Irvine, L. Sun, L. Ma, P.D. Weyman, D.G. Gibson, J.I. Glass, J.C. Venter, C.A. Hutchison, H.O. Smith, Y. Suzuki, Transferring whole genomes from bacteria to yeast spheroplasts using entire bacterial cells to reduce DNA shearing, Nat. Protoc. 9 (2014) 743–750. 10.1038/nprot.2014.045.

[26] M.P.M. Soltysiak, R.S. Meaney, S. Hamadache, P. Janakirama, D.R. Edgell, B.J. Karas, Trans-kingdom conjugation within solid media from Escherichia coli to Saccharomyces cerevisiae, Int. J. Mol. Sci. 2019, Vol. 20, Page 5212. 20 (2019) 5212. 10.3390/IJMS20205212.

[27] M. Kostylev, A.E. Otwell, R.E. Richardson, Y. Suzuki, Cloning Should Be Simple: Escherichia coli DH5α-Mediated Assembly of Multiple DNA Fragments with Short End Homologies, PLoS One. 10 (2015) e0137466. 10.1371/journal.pone.0137466.

[28] B.J. Karas, C. Tagwerker, I.T. Yonemoto, C.A. Hutchison, H.O. Smith, Cloning the Acholeplasma laidlawii PG-8A Genome in Saccharomyces cerevisiae as a Yeast Centromeric Plasmid, ACS Synth. Biol. 1 (2012) 22–28. 10.1021/sb200013j.

