## Supplemental file for "Deletion of HindIIR and HindIIIR improves DNA transfer via electroporation to *Haemophilus influenzae* Rd"

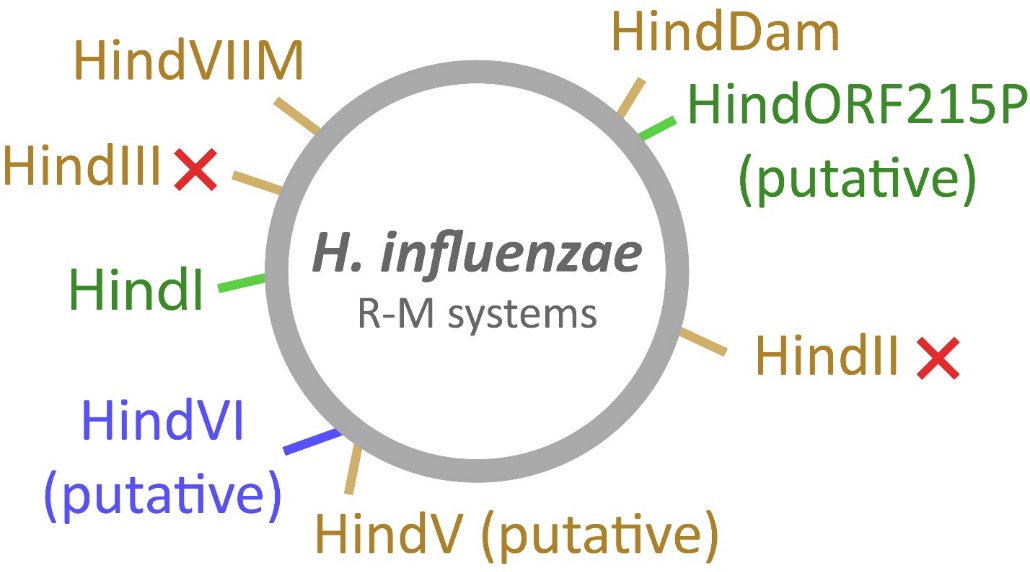

**Figure S1. Map of the *H. influenzae* chromosome with its RM systems labeled.** HindVI and HindV methylases have been experimentally validated whereas the respective endonucleases have not; HindORF215P has not been experimentally tested. Green, brown, and blue labels correspond to type I, type II, and type III RM systems, respectively. Red X’s mark the RM systems that have been removed in ΔHindII/III [1]. Figure adapted from REBASE [2].

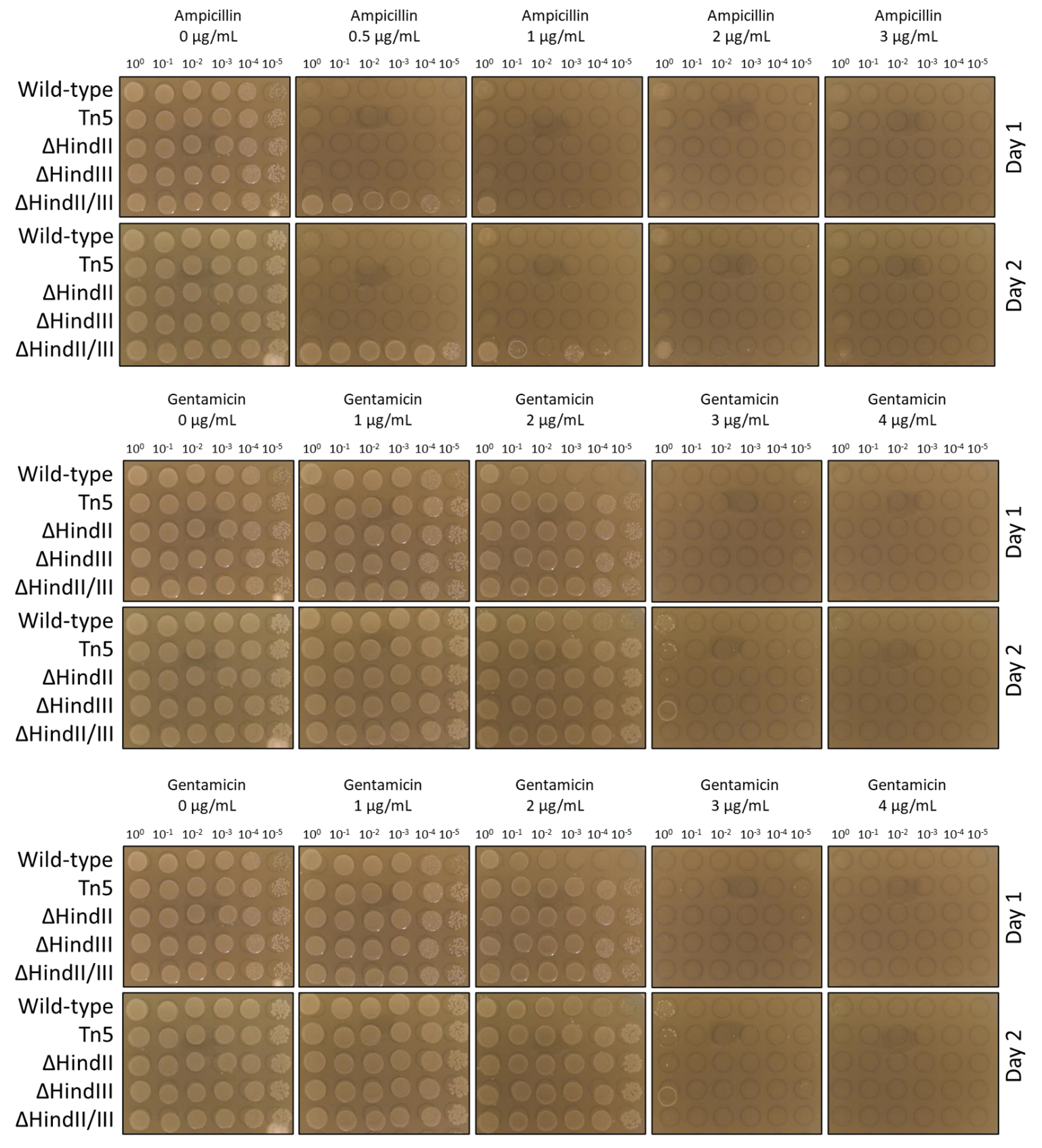

**Figure S2. Sensitivity of *H. influenzae* strains to various antibiotics.**

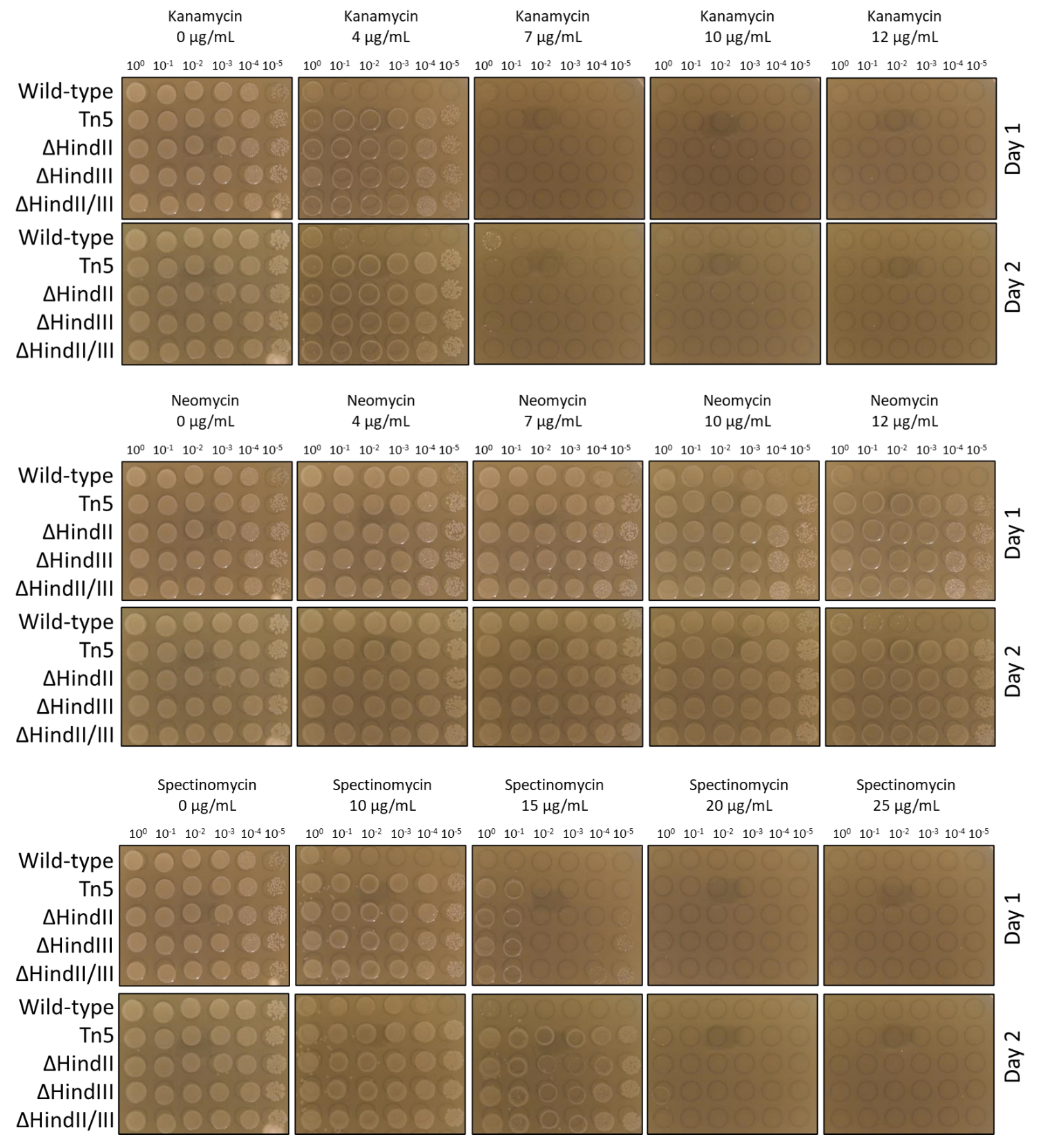

**Figure S2. Sensitivity of *H. influenzae* strains to various antibiotics.**

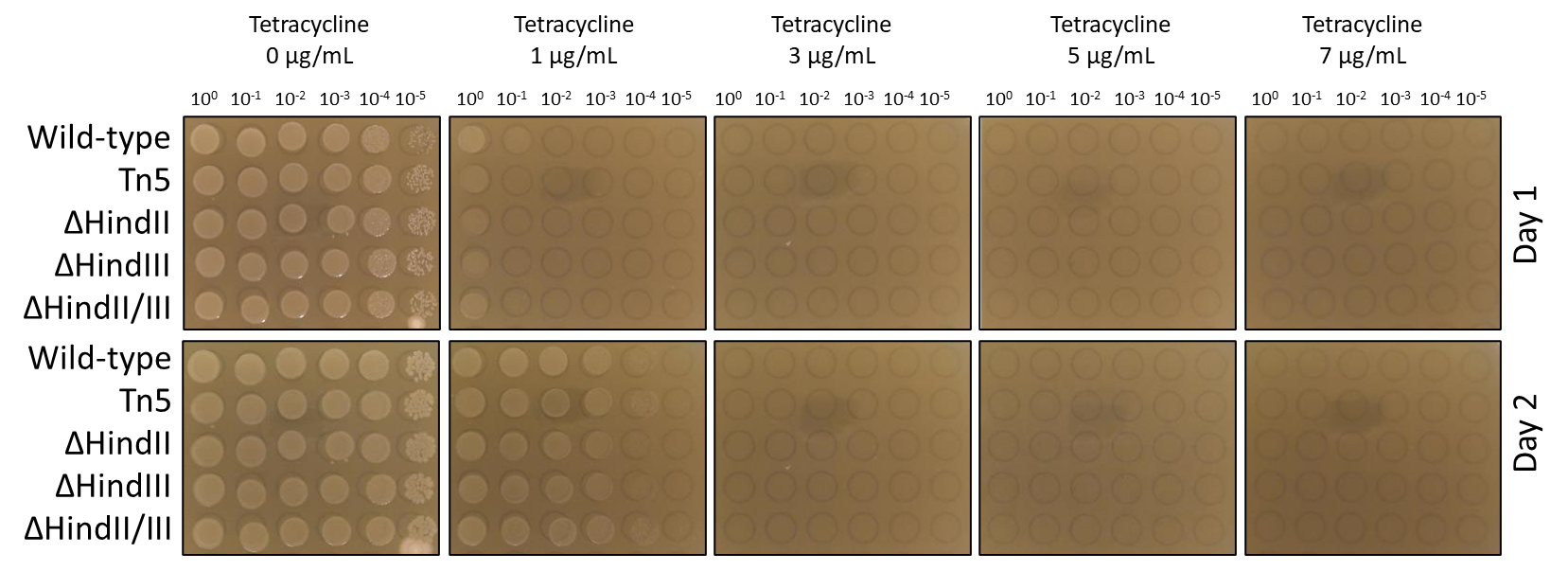

**Figure S2. Sensitivity of *H. influenzae* strains to various antibiotics.** Wild-type *H. influenzaae* Rd KW20 as well as four strains generated by Karas et al. 2013 were evaluated for sensitivity to antibiotics at different strengths. “Tn5” corresponds to a strain of *H. influenzae* where a Tn5 transposon carrying a chloramphenicol resistance marker and yeast vector was inserted at a random location. “ΔHindII” and “ΔHindIII,” the same yeast vector and chloramphenicol resistance marker were used to replace the corresponding restriction endonuclease gene. “ΔHindII/III” was generated by taking ΔHindIII and adding a cassette containing tetracycline and ampicillin resistance markers to replace the ΔHindII gene. To assess the sensitivity of each of these strains to antibiotics, cultures were grown overnight and diluted to an OD_600_ ≈ 0.1, then serially diluted before spot-plating onto sBHI plates containing varying concentrations of each antibiotic.

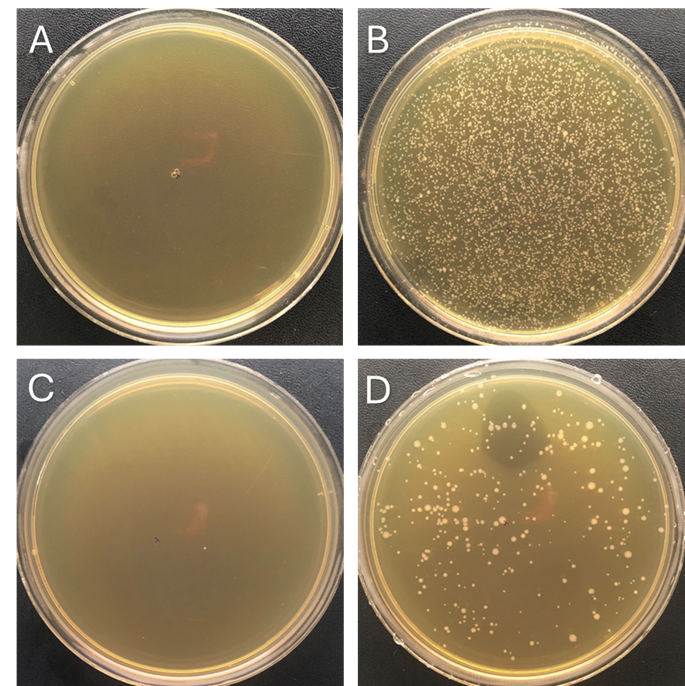

**Figure S3. Selection of transformants with tetracycline and kanamycin.** (**A**) 100 µL of negative control plated on sBHI supplemented with 5 µg/mL of tetracycline. (**B**) 100 µL of pHflu8 transformation plated on 5 µg/mL tetracycline. (**C**) 10 µL of negative control plated on 20 µg/mL kanamycin. (**D**) 10 µL of pHflu3 transformation plated on 20 µg/mL kanamycin.

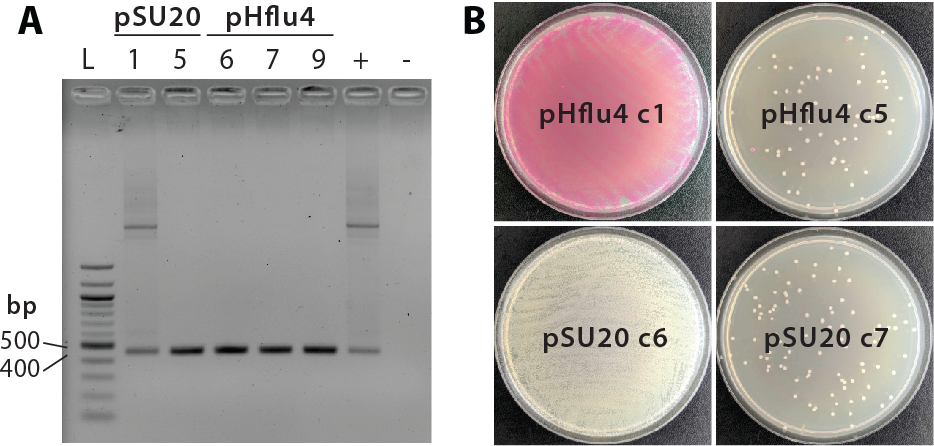

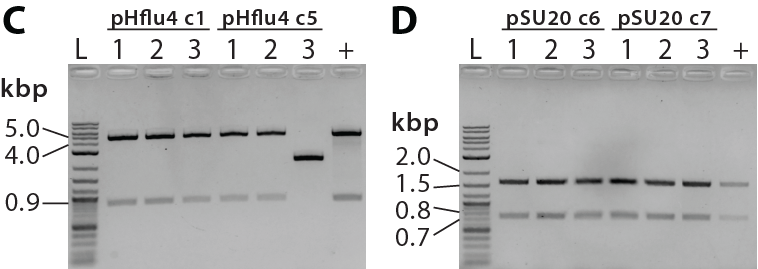

**Figure S4. Validation of plasmids isolated from wild-type *H. influenzae* transformant colonies.** (**A**) DNA isolated from wild-type *H. influenzae* transformants was screened by PCR for the presence of the chloramphenicol acetyltransferase (*cat*) gene. Lane numbers correspond to those from **Figure 1B**. All colonies screened reveal the presence of *cat*, however it is possible that the gene was integrated into the genome rather than maintained on the appropriate plasmid. +: pSU20 isolated from *E. coli* as a positive control; -: water instead of PCR template as negative control; L: NEB 100-bp ladder. (**B**) DNA from wild-type *H. influenzae* transformants was used to transform *E. coli* by electroporation. DNA from *H. influenzae* transformants showing the correct restriction digest pattern as seen in Figure 1B produced significantly more *E. coli* transformants, suggesting that the correct plasmid was not present in the transforming DNA. (**C-D**) DNA, isolated from individual *E. coli* colonies from the plates shown in B-E, was digested by EcoRI (pHflu4) or ApoI (pSU20) to screen for plasmid DNA. Overall, this analysis indicates that not only are lower numbers of transformants obtained with the wild-type strain, but also some of the transformed plasmids will have mutations that make the wild-type strain less suitable for synthetic biology applications.

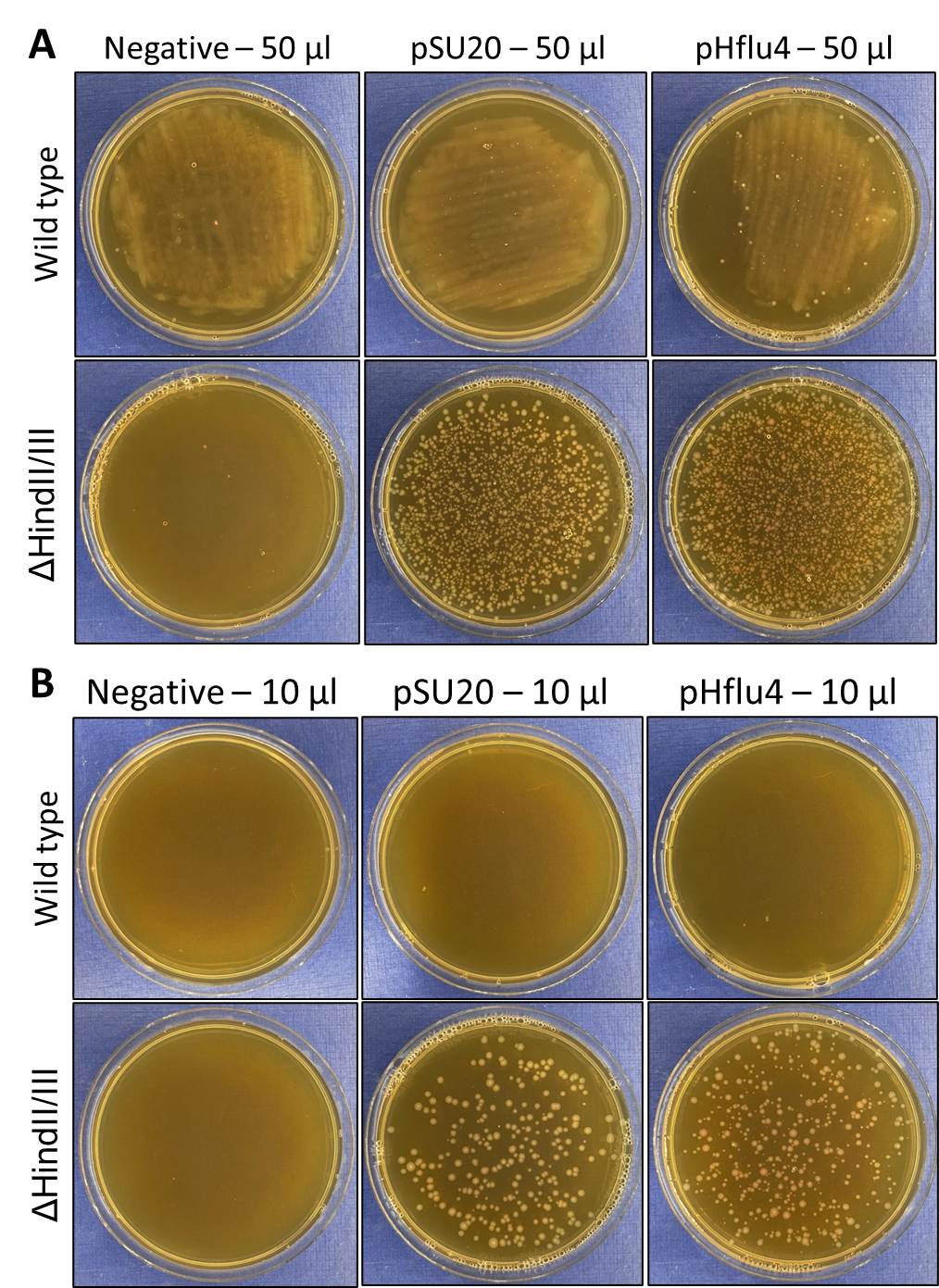

**Figure S5. Background growth of wild-type *H. influenzae* but not those of ΔHindII/III.** **(A)** 50 µL of electroporated wild-type *H. influenzae* Rd KW20 and ΔHindII/III cells were plated on sBHI plates supplemented with 3 µg/mL of chloramphenicol. Note that background growth/biofilm is visible on all wild type plates: negative and cells transformed with pSU20 or pHflu4 plasmids. **A)** 10 µL of electroporated wild-type *H. influenzae* Rd KW20 and ΔHindII/III cells were plated on sBHI plates supplemented with 3 µg/mL of chloramphenicol. Note that with this amount of cells no background growth/biofilm is visible for wild type.

**Table S1. List of primers used in this study.**

| **Name** | **Sequence** |
| --- | --- |
| SH01 | GGAGCACCTCAAAAACACCATCATACACTAAATCATTAATTAATAAGAGCTTGGTGAGCG |
| SH02 | CACAGGTATTTATTCGGCGCAAAGTGCGTCGGGTGATGCTAGCTCGATCGTCTTGCCTTG |
| SH03 | GTAAGTACATCACCGACGAGCAAGGCAAGACGATCGAGCTAGCATCACCCGACGCACTTT |
| SH04 | CTCCTAGCGCTCACCAAGCTCTTATTAATTAATGATTTAGTGTATGATGGTGTTTTTGAG |
| SH05 | CCGGCGGTGCTTTTGCC |
| SH06 | GAGTCATCCGCTAGGTGGA |
| SH07 | ACTATGAGCACGTCCGCGA |
| SH08 | TACGCCCGGTAGTGATCTTATT |
| SH09 | GGGTACCGAGCTCGAATTCACTGGCCGTCGTTTTACAACGTCGTGACTGGGAAAACCCTG |
| SH10 | CCTCGAGGGGCATGCAAGCTTGGCGTAATCATGGTCATAGCTGTTTCCTGTGTGAAATTG |
| SH11 | ACCATGATTACGCCAAGCTTGCATGCCCCTCGAGGCGCGGAACCCCTATTTGTTTATTTT |
| SH12 | GACGTTGTAAAACGACGGCCAGTGAATTCGAGCTCGGTACCCTACGGGGTCTGACGCTCA |
| SH13 | AGCTATGACCATGATTACGCCAAGCTTGCATGCCCCTCGAGGTCGAGCGACAGGGCGAAG |
| SH14 | CGTTGTAAAACGACGGCCAGTGAATTCGAGCTCGGTACCCCAGGGTTATGCAGCGGAAGA |
| SH15 | TCACACTGGCTCACCTTCGGGTGGGCCTTTCTGCGTTTATAGGGTACCGAGCTCGAATTC |
| SH16 | TCTGTGAGCTAGCATTATACCTAGGACTGAGCTAGCTGTCAACCTCGAGGGGCATGCAAG |
| SH17 | ACCATGATTACGCCAAGCTTGCATGCCCCTCGAGGTTGACAGCTAGCTCAGTCCTAGGTA |
| SH18 | TTGTAAAACGACGGCCAGTGAATTCGAGCTCGGTACCCTATAAACGCAGAAAGGCCCACC |
| SH19 | CAGCTATGACCATGATTACGCCAAGCTTGCATGCCCCTCGAGGGTAAAACGACGGCCAGT |
| SH20 | CGACGTTGTAAAACGACGGCCAGTGAATTCGAGCTCGGTACCCCAGGAAACAGCTATGAC |
| SH21 | CCATGATTACGCCAAGCTTGCATGCCCCTCGAGGTCCCTTTAGTGAGGGTTAATGTCGTG |
| SH22 | AATTCGAGCTCGGTACCCTTATGTGATTTTGTTGAACATATAACGAACTTTATCAATACG |

**References:**

[1]	B.J. Karas, J. Jablanovic, L. Sun, L. Ma, G.M. Goldgof, J. Stam, A. Ramon, M.J. Manary, E.A. Winzeler, J.C. Venter, P.D. Weyman, D.G. Gibson, J.I. Glass, C.A. Hutchison, H.O. Smith, Y. Suzuki, Y. Suzuki, Direct transfer of whole genomes from bacteria to yeast., Nat. Methods. 10 (2013) 410–412. https://doi.org/10.1038/nmeth.2433.

[2]	R.J. Roberts, T. Vincze, J. Posfai, D. Macelis, REBASE: a database for DNA restriction and modification: enzymes, genes and genomes, Nucleic Acids Res. 51 (2023). https://doi.org/10.1093/nar/gkac975.
